# Genetic and environmental contributions to ancestry differences in gene expression in the human brain

**DOI:** 10.1101/2023.03.28.534458

**Authors:** Kynon J.M. Benjamin, Qiang Chen, Nicholas J. Eagles, Louise A. Huuki-Myers, Leonardo Collado-Torres, Joshua M. Stolz, Geo Pertea, Joo Heon Shin, Apuã C.M. Paquola, Thomas M. Hyde, Joel E. Kleinman, Andrew E. Jaffe, Shizhong Han, Daniel R. Weinberger

**Affiliations:** Lieber Institute for Brain Development, Baltimore, MD, USA; Department of Neurology, Johns Hopkins University School of Medicine, Baltimore, MD, USA; Department of Psychiatry and Behavioral Sciences, Johns Hopkins University School of Medicine, Baltimore, MD, USA; Center for Computational Biology, Johns Hopkins University, Baltimore, MD, USA; Department of Neuroscience, Johns Hopkins University School of Medicine, Baltimore, MD, USA; Neumora Therapeutics, Watertown, MA, USA; Department of Genetic Medicine, Johns Hopkins University School of Medicine, Baltimore, MD, USA

## Abstract

Ancestral differences in genomic variation are determining factors in gene regulation; however, most gene expression studies have been limited to European ancestry samples or adjusted for ancestry to identify ancestry-independent associations. We instead examined the impact of genetic ancestry on gene expression and DNA methylation (DNAm) in admixed African/Black American neurotypical individuals to untangle effects of genetic and environmental factors. Ancestry-associated differentially expressed genes (DEGs), transcripts, and gene networks, while notably not implicating neurons, are enriched for genes related to immune response and vascular tissue and explain up to 26% of heritability for ischemic stroke, 27% of heritability for Parkinson’s disease, and 30% of heritability for Alzhemier’s disease. Ancestry-associated DEGs also show general enrichment for heritability of diverse immune-related traits but depletion for psychiatric-related traits. The cell-type enrichments and direction of effects vary by brain region. These DEGs are less evolutionarily constrained and are largely explained by genetic variations; roughly 15% are predicted by DNAm variation implicating environmental exposures. We also compared Black and White Americans, confirming most of these ancestry-associated DEGs. Our results highlight how environment and genetic background affect genetic ancestry differences in gene expression in the human brain and affect risk for brain illness.

**Summary:** We examine the impact of genetic ancestry on gene expression and DNA methylation of admixed African/Black Americans, highlighting how genetic and environmental background affect risk for brain illness.

## Introduction

Health disparities have endured for centuries (*1*). In neuroscience and genomics, individuals with recent African genetic ancestry (AA) account for less than 5% of large-scale research cohorts for brain disorders but are 20% more likely to experience a major mental health crisis (*2*, *3*). Insights gained from genome-wide association studies (GWAS) about disease risk are promising for clinical applications (e.g., drug targets for novel therapeutics and polygenic risk prediction). However, the majority of GWAS of brain-related illness lack diversity with regards to inclusion of AA individuals, who account for less than 5% of GWAS participants (*4*), despite AA individuals having more extensive genetic variation than any other population. This lack of diversity limits the accuracy of genetic risk prediction, hinders the development of effective personalized neurotherapeutics for non-European genetic ancestry (EA) individuals (*5*), and limits our potential for novel discovery. While diversity in large-scale GWAS has increased in recent years (e.g., 1000 Genomes Project (*6*), All of Us research program, Trans-Omics for Precision Medicine [TOPMed] (*7*), and Human Heredity and Health in Africa [H3Africa] Consortium (*8*)), population-based genetic association studies do not directly elucidate potential biological mechanisms of risk variants.

To bridge this gap, we need studies of the biological impact of genetic variation on molecular traits (e.g., mRNA and DNA methylation [DNAm]) in disease-relevant tissues of diverse populations. Recent efforts to bridge this gap with cross-ancestry expression quantitative trait loci (eQTL) have focused on improved fine mapping while leaving unanswered the question of how gene expression and epigenetic regulation are parsed specifically by ancestry (*9*). Despite a clear urgent need, no large-scale studies examine the biological impact of genetic ancestry on gene expression in the human brain focused on the differences between AA and EA.

An obvious impediment to undertaking this task is the limited availability of brain tissue from AA individuals. Currently, the most widely used resource for human postmortem tissue is the Gene-Tissue Expression Project (GTEx), which has publicly available RNA-sequencing and single nucleotide polymorphisms (SNP) genotyping for nearly 1,000 mostly elderly individuals, including data from 13 brain regions (114 to 209 individuals per region). However, the majority of GTEx brain samples are of EA, and for some brain regions, GTEx has no non-EA individuals. In comparison, the BrainSeq Consortium, a collaboration between seven pharmaceutical companies and the Lieber Institute for Brain Development (LIBD), has one of the largest postmortem brain collections of psychiatric disorders, including 784 Black American samples across 587 unique individuals, with a mean age of 44. While reports from this consortium and other large-scale analyses in the brain – including from the hippocampus, caudate nucleus (“caudate”), dorsolateral prefrontal cortex (DLPFC), and granule cells of the dentate gyrus (“dentate gyrus”) – have samples of diverse genetic ancestry (*10–16*), they have typically been “adjusted” for ancestry status, which limits our understanding of ancestry-specific effects in the brain.

To address these gaps, here we use the LIBD RNA-sequencing, SNP genotype, and whole genome bisulfite sequencing (WGBS) datasets to evaluate genetic and environmental contributions to genetic ancestry differences in gene expression in the human brain (**Fig. 1**). We identify transcriptional features associated with genetic ancestry (African or European) in admixed neurotypical Black American donors (n=151). We quantify the contributions of common genetic variations to genetic ancestry differences using a total of 425 samples, including the caudate (n=122), dentate gyrus (n=47), DLPFC (n=123), and hippocampus (n=133). Additionally, we examine the influence of genetic ancestry on DNAm using WGBS data of the admixed Black American donors from the caudate (n=89), DLPFC (n=69), and hippocampus (n=69). To confirm the genetic ancestry-associated differences in gene expression and to highlight the effect of environment by genetic ancestry differences, we further examine transcriptional and DNAm differences in individuals of limited admixture (Black Americans ≥ 0.8 AA and White Americans > 0.99 EA).

**Fig. 1:**
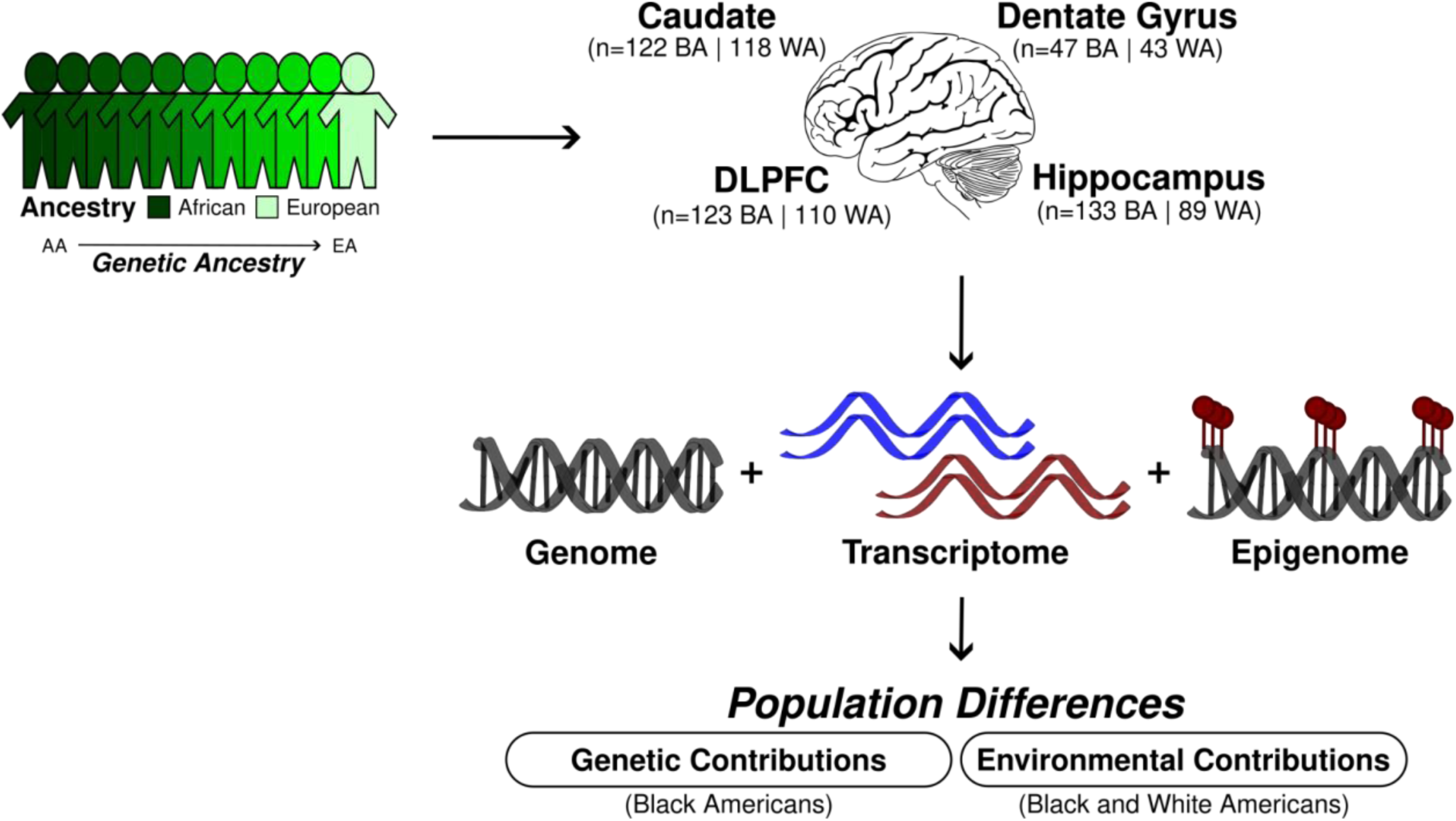
Study design for the examination of the genetic and environmental contributions to genetic ancestry-associated expression differences. BA stands for Black Americans and WA for White Americans.

## Results

### Significant enrichment of immune response for differential expression associated with genetic ancestry across the brain

We selectively examined our admixed Black American population (151 unique individuals; **Table S1**) to 1) characterize transcriptional changes associated with African or European genetic ancestry in neurotypical adults (age > 17) and 2) limit potential confounding effects of systematic environmental factors that may differ between Black and White American samples. These analyses included RNA sequencing data from caudate (n=122), dentate gyrus (n=47), DLPFC (n=123), and hippocampus (n=133). Our admixed Black American population showed a varied proportion of EA (STRUCTURE (*17*); EA mean = 0.21, range = 0-0.62; **Fig. S1**) consistent with previous reports (*18*, *19*). As such, we used these continuous genetic ancestry estimates to identify differentially expressed features (genes, transcripts, exons, and junctions) that were linearly correlated with ancestry levels and adjusted for sex, age, and RNA quality. This RNA quality adjustment includes experiment-based RNA degradation metrics (obtained with the qSVA methodology) that account for batch effect and cell composition (*12*, *20*). To increase our power of detection and improve effect size estimates, we applied the multivariate adaptive shrinkage (“mash” (*21*)) method, which leverages the correlation structure of genetic ancestry effects across brain regions (see **Methods** for details). Of the 16,820 genes tested, we identified 2,570 (15%; 1,437 of which are protein coding) unique differentially expressed genes (DEGs) based on ancestry variation (local false sign rate [lfsr] < 0.05; **Fig. 2A**, **Table S2**, and **Data S1**) across the caudate (n=1,273 DEGs), dentate gyrus (n=997), DLPFC (n=1,075), and hippocampus (n=1,025). While this number increased when we examined differential expression based on local ancestry (9,906 [62% of genes tested]; 6,982 protein coding; **Table S3**) across the caudate (n=6,657 DEGs), dentate gyrus (n=4,154), DLPFC (n=6,148), and hippocampus (n=7,006), effect sizes between global- and local-ancestry DEGs showed significant positive correlations (all Spearman; rho > 0.57, p-value < 0.01; **Fig. S3**) across all brain regions. When examining isoform-level associations (transcripts, exons, and junctions), we found an additional 8,012 unique global ancestry-associated DEGs (lfsr < 0.05; **Fig. S2**, **Table S2**, and **Data S1**) and 6,629 unique local ancestry-associated DEGs (lfsr < 0.05; **Table S3** and **Data S2**) in these Black Americans. Similarly, we found that isoform-level local ancestry DE features showed significant positive correlation in effect sizes compared with global ancestry DE features (**Fig. S3**).

**Fig. 2:**
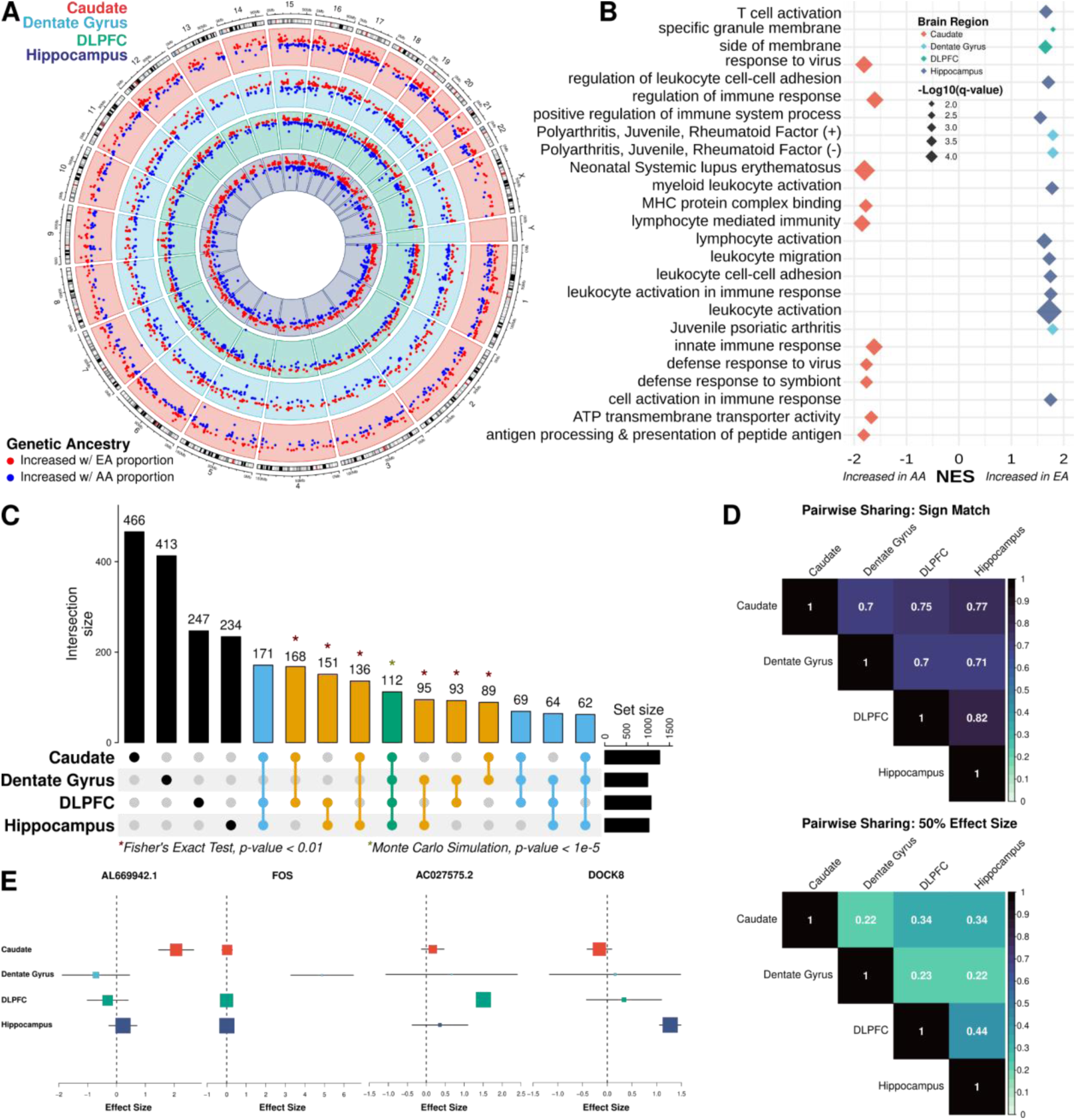
Extensive ancestry-associated expression changes across the brain region. **A.** Circos plot showing ancestry DEGs across the caudate (red), dentate gyrus (blue), DLPFC (green), and hippocampus (purple). **B.** Gene set enrichment analysis (GSEA) of differential expression analysis across brain regions, highlighting terms associated with increased AA (African ancestry) or EA (European ancestry) proportions. **C.** UpSet plot showing large overlap between brain regions. Green is shared across the four brain regions; blue, shared across three brain regions; orange, shared between two brain regions; and black, unique to a specific brain region. * Indicating significant pairwise enrichment (Fisher’s exact test) or significant overlap between all four brain regions (Monte Carlo simulation). **D.** Heatmaps of the proportion of ancestry DEG sharing with concordant direction (sign match; top) and within a factor 0.5 effect size (bottom) **E.** Metaplot showing examples of brain region-specific ancestry effects.

To evaluate the functional aspects of these genetic ancestry-associated DEGs (global and local ancestry), we performed gene set enrichment analysis with the Gene Ontology (GO) and Disease Gene Network (DisGeNET (*22*)) databases for each brain region. It is noteworthy that while there was no enrichment of neuronal gene sets, we observed significant enrichment (GSEA and hypergeometric, q-value < 0.05) for GO and DisGeNET terms primarily related to immune response, including innate, adaptive, and virus responses (**Data S3, Fig. 2B**, and **Fig. S4**). Interestingly, the caudate showed an opposite direction of effect compared with the DLPFC and hippocampus. Specifically, the caudate showed enrichment of immune response associated with DEGs upregulated in relation to AA proportion, while dentate gyrus, DLPFC, and hippocampus showed enrichment for immune-related pathways associated with DEGs upregulated in EA proportion (**Fig. 2B** and **Fig. S5**). While not significant, we observed the same pattern of opposite directionality of effect for immune-related pathways with local ancestry-associated DEGs (**Fig. S6**).

When we expanded this analysis to the isoform level (transcripts, exons, and junctions), we also found significant association with immune-related pathways and similar directions of effect (upregulated for AA proportion in the caudate and upregulated for EA proportion in dentate gyrus, DLPFC, and hippocampus). Furthermore, we also found significant analogous enrichment of these DEGs for genes with population differences in macrophages (*18*) associated with innate immune response to infection (Fisher’s exact test, false discovery rate [FDR] < 0.05; **Fig. S7**). Additionally, we found significant enrichment (Fisher’s exact test, FDR < 0.01) for ancestry-associated DEGs (global ancestry) in gene coexpression network modules generated using WGCNA (Weighted Gene Co-expression Network Analysis (*23*); **Fig. S8**). Consistent with our DEG analysis, the immune response pathway enrichment in these modules showed analogous opposite direction of effects based on region (**Fig. S9**).

Observing an enrichment of the immune response pathway in bulk tissue, we performed cell-type (*24*, *25*) enrichment analysis to evaluate the cellular context of these ancestry-associated DEGs (global ancestry). We found significant enrichment (Fisher’s exact test, FDR < 0.05; **Fig. S10** and **Fig. S11A**) for genes specifically expressed in brain immune cells (i.e., glia and microglia cell types) and neurovasculature (i.e., pericyte, endothelial, and vascular tissue cells), but not peripheral immune cells. Additionally, we observed enrichment for distinct subtypes of glial cells (*26*) (**Fig. S12**). Interestingly, local ancestry-associated DEGs showed significant enrichment for brain and non-brain immune cells (Fisher’s exact test, FDR < 0.05; **Fig. S13** and **Fig. S11B**) potentially due to the larger number of detected DEGs. Even so, we found the level of enrichment of non-brain immune cells (global and local) on average smaller than brain immune cells. Remarkably, we again found primarily significant depletion of DEGs (global and local) for any genes specific to neuronal cell types. Consistently, we observed those immune-related pathways and associated cell types (i.e., microglia and perivascular macrophage) for DEGs upregulated with increasing AA proportion in the caudate and upregulated with increasing EA proportion in the dentate gyrus, DLPFC, and hippocampus. Although we found some glial cell subtype (*26*) composition differences (ANOVA, FDR < 0.05; **Fig. S14**) using publically available single cell data from brain regions with similar composition (*27*), no glial subtype (*26*) showed specificity for a specific direction of ancestry effect (**Fig. S12**). Altogether, these results suggest that ancestry-associated DEGs in the human brain are strongly associated with the brain-specific immune response, and specific direction of effects vary according to brain region.

### Sharing of genetic ancestry-associated expression differences across the brain

To understand the regional specificity of global ancestry-associated differentially expressed features, we compared DEGs from each brain region and observed extensive sharing across regions. Specifically, we observed 1,210 DEGs (47.1%) shared between at least two brain regions, where all pairwise overlaps demonstrated significant enrichment (Fisher’s exact test, p-value < 0.01; **Fig. 2C**). Moreover, 478 DEGs (18.6%) were shared among at least three brain regions with a significant overlap of 112 of these DEGs (4.4%; Monte Carlo simulation, p-value < 1e-5) across all four brain regions.

Interestingly, 27 of the 112 shared DEGs (24%) showed discordant direction of effect in at least one of the four brain regions. This correlated well with the pairwise correlation of shared DEGs that shared direction of effect (70% to 82%; **Fig. 2D**). While shared direction of effect across brain regions was relatively high, this proportion of sharing dropped substantially when effect size was taken into account (0.22 to 0.44; **Fig. 2D**). Corresponding with the large proportion of discordant DEGs, we also found a large number of brain region-specific DEGs (1360 [52.9%]; **Fig. 2E**), which increased when considering isoform-level analysis (transcript [63.6%], exon [67.6%], and junction [69.7%]). This is consistent with other studies that show isoform-level brain region specificity (*28*).

### HLA region and immune cell composition play a limited role on ancestry-associated expression differences across the brain

Given the primary enrichment signal for immune-related pathways and cell types, we next investigated if immune variation was driving the observed transcriptional changes. Initially, we examined enrichment of ancestry-associated DEGs for the major histocompatibility complex (MHC) region. Here, we found global ancestry-associated DEGs of the caudate, DLPFC, and hippocampus enriched for HLA class II, while dentate gyrus enriched for Zinc finger proteins associated with the extended class I MHC region (Fisher’s exact test, FDR < 0.05; **Fig. S15**). While we found limited enrichment of local ancestry-associated DEGs for gene clusters of the MHC region across brain regions, we still observed significant enrichment of HLA class II genes for the caudate similar to global ancestry DEGs (Fisher’s exact test, FDR < 0.05; **Fig. S16**). Altogether, these results suggest that ancestry-associated DEGs (global and local) within the MHC region are primarily enriched for HLA class II genes.

Next, we re-examined functional enrichment of ancestry-associated DEGs after removing the MHC region (i.e., HLA-specific genes, MHC region, and extended MHC region) to determine if the limited MHC enrichment drove functional enrichment of immune-related pathways of our ancestry-associated DEGs. After excluding the HLA genes, we still observed strong enrichment for immune-related pathways (**Fig. S17**). Furthermore, we observed similar immune-related enrichment (i.e., response to virus, interleukin-12 production, macrophage activation, leukocyte migration, and innate immune response) after excluding the MHC region (**Fig. S18**) or the extended MHC region (**Fig. S19**) across brain regions. This was also the case with local ancestry DEGs (**Fig. S20**), suggesting that the extended MHC region does not drive ancestry-associated DEG enrichment of immune-related pathways.

Although the MHC region did not appear to drive our immune response enrichment, immune variation either from HLA gene diversity or glial cell composition could still contribute to the transcriptional changes observed in our ancestry-associated DEGs. As such, we next assessed to what degree HLA variation or glial cell composition contributed to the expression changes. To assess glial cell composition, we added glial cells composition (astrocytes, microglia, macrophage, oligodendrocytes, oligodendrocyte progenitor cells, and T cells) as covariates in our DE model. When we compared effect sizes with the original model, we found a high degree of correlation (Spearman; rho from 0.81 to 0.92; **Fig. S21A**), suggesting glial cell composition had a minimal effect. For HLA variation, we added the first five PCs of imputed HLA alleles (accounting for 66% of variance explained) as covariates and compared effect sizes with our original model. Similar to glial cell composition, we found HLA genetic variation only minimally changed effect sizes (Spearman; rho from 0.83 to 0.87; **Fig. S21B**). Altogether, these sensitivity analyses suggest that immune variation contributes only minimally to transcriptional changes for ancestry-associated DEGs.

### Ancestry-associated DEGs are evolutionarily less constrained

With consistent significant enrichment of DEGs and co-expression modules for immune response, we hypothesized that this functional connection for the DEGs with a cellular biology that is uniquely adaptable would render them more likely to be tolerant of phenotypic consequences of gene disruption and would therefore be evolutionarily less constrained. To test this hypothesis, we examined the gene and transcript constraint scores (*29*) for the global ancestry-associated DEGs. Unsurprisingly, we found significant depletion of DEGs for highly constrained genes (Fisher’s exact test, FDR < 0.0001; **Fig. 3A**). On the transcript level, we found a similar trend (**Fig. 3B**) with differentially expressed (DE) transcripts associated with less constrained genes. Additionally, we observed a significant negative correlation with DEGs signal (lfsr) and gene and transcript constraint scores (Pearson, p-value < 0.0001; **Fig. 3C**). Unsurprisingly, these results suggest that ancestry-associated DE features are associated with the more rapidly evolving genes as previously seen in immunity related genes (*30*, *31*).

**Fig. 3:**
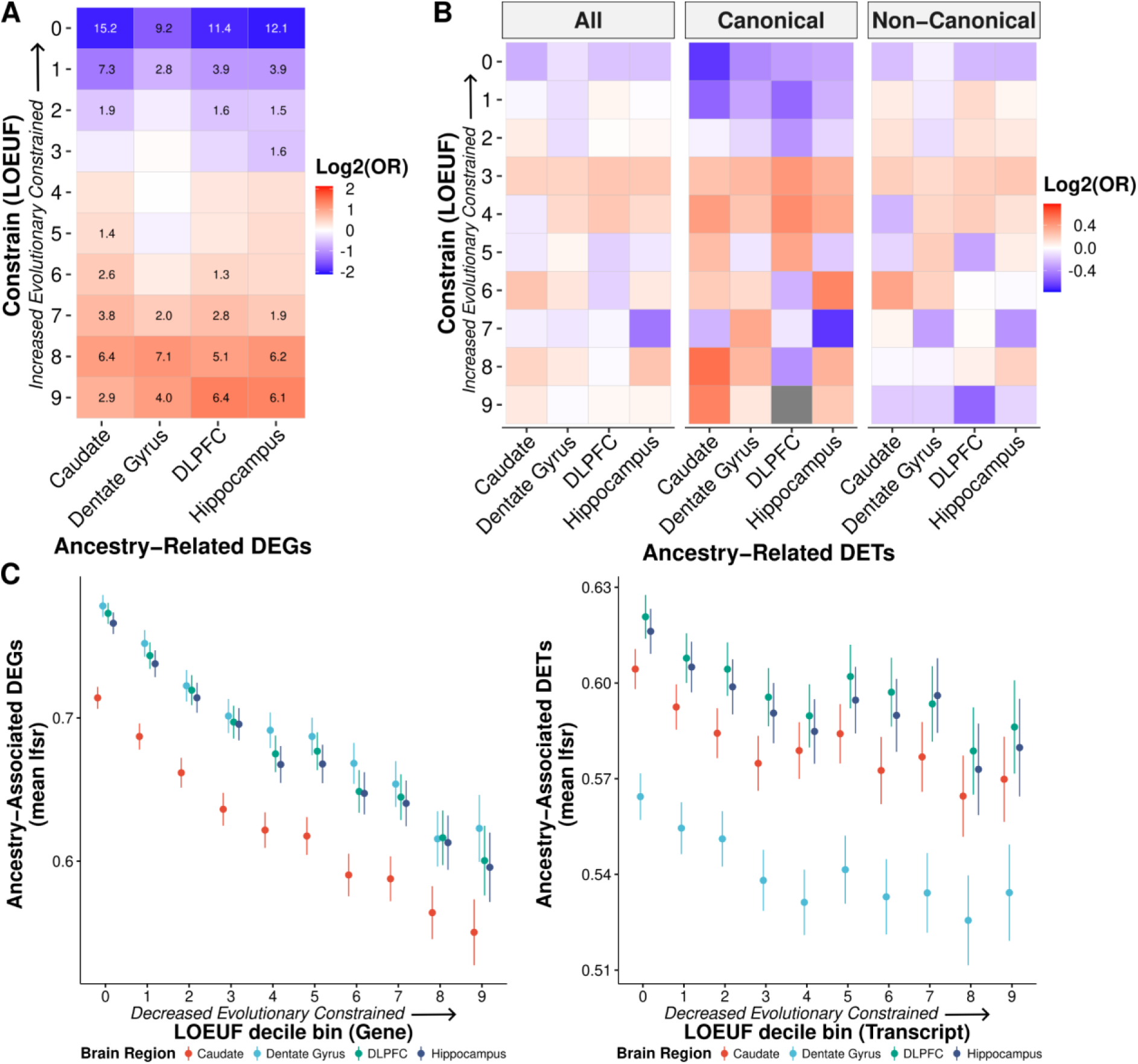
Ancestry-associated genes and canonical transcripts are evolutionarily less constrained. **A.** Significant depletion of ancestry DEGs for evolutionarily constrained genes (canonical transcripts) across brain regions. Significant depletion/enrichments (two-sided, Fisher’s exact test, FDR corrected p-values, −log10 transformed) are annotated within tiles. Odds ratios (OR) are log_2_ transformed to highlight depletion (blue) and enrichment (red). **B.** Similar trend of depletion of ancestry DE transcripts (DETs; all, canonical, and non-canonical) for evolutionarily constrained transcripts across brain regions. Odds ratios are log_2_ transformed to highlight depletion (blue) and enrichment (red). **C.** The mean of ancestry-associated DE feature (i.e., gene and transcript) lfsr as a function of LOEUF (loss-of-function observed/expected upper bound fraction) decile shows a significant negative correlation for genes (left; for the caudate, dentate gyrus, DLPFC, and hippocampus: two-sided, Pearson, *r* = −0.20, −0.20, −0.21, and −0.21; p-value = 3.0×10^−122^, 7.6 × 10^−113^, 8.6×10^−126^, and 1.2 × 10^−122^) and transcripts (right; for the caudate, dentate gyrus, DLPFC, and hippocampus: two-sided, Pearson, *r* = −0.05, −0.05, −0.04, and −0.04; p-value = 8.6×10^−13^, 1.7×10^−11^, 9.0×10^−11^, and 3.2 × 10^−10^). Error bars correspond to 95% confidence intervals.

### The role of genetic variants on ancestry-associated expression differences in the brain

To assess the contribution of genetic variation to genetic ancestry-associated DEGs, we first mapped main effect cis-eQTL in Black American individuals (n=120, 45, 121, and 131 for the caudate, dentate gyrus, DLPFC, and hippocampus, respectively) examining genetic variants within +/-500 kb of each feature (gene, transcript, exon, and junction). To improve detection of eQTL, we applied mash and identified at least one cis-eQTL for 13,857 genes (“eGenes”) across brain regions (lfsr < 0.05; n=10,867 for the caudate; n=11,664 for the dentate gyrus; n=11,173 for the DLPFC; and n=10,408 for the hippocampus; **Table S4** and **Data S6**). Of these 13,857 eGenes, the majority (64.1%; **Fig. 4A**) were shared across all brain regions with only about 0.25 to 14.5% showing brain region specificity. When we examined the direction of effect, however, this number dramatically increased with more than 96% sign matching (**Fig. 4B**).

**Fig. 4:**
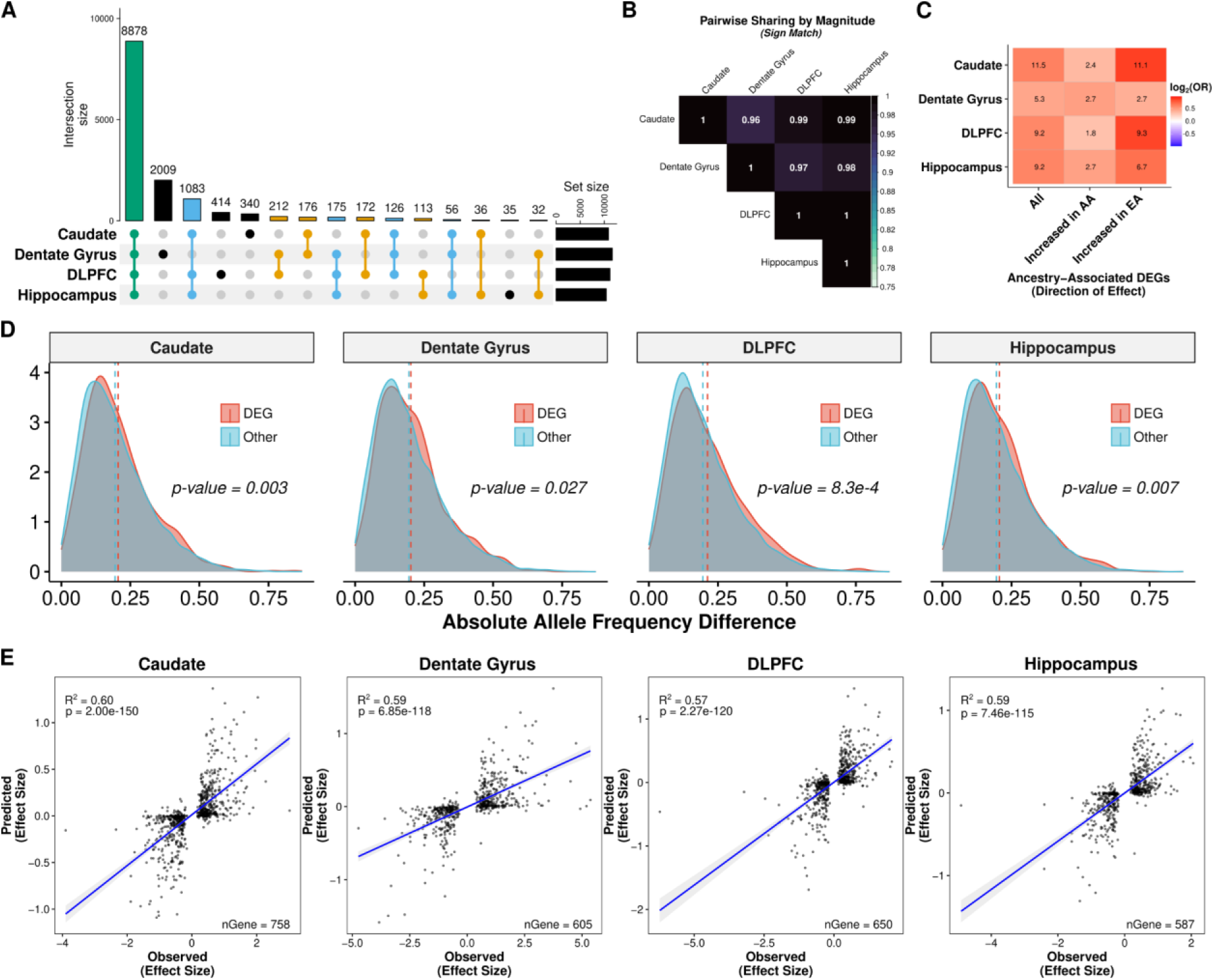
Genetic contribution of genetic ancestry differences in expression across the brain. **A.** UpSet plot showing large overlap between brain regions of eGenes. **B.** Heatmap of the proportion of ancestry DEG sharing with concordant direction (sign match). **C.** Significant enrichment of ancestry-associated DE genes for eGenes (unique gene associated with an eQTL) across brain regions separated by direction of effect (increased in AA or EA proportion). **D.** Density plot showing significant increase in absolute allele frequency differences (AFD; one-sided, Mann-Whitney U, p-value < 0.05) for global ancestry-associated DEGs (red) compared with non-DEGs (blue) across brain regions. A dashed line marks the mean absolute AFD. Absolute AFD calculated as the average absolute AFD across a gene using significant eQTL (lfsr < 0.05). **E.** Correlation (two-sided, Spearman) of elastic net predicted (y-axis) versus observed (x-axis) ancestry-associated differences in expression among ancestry-associated DEGs with an eQTL across brain regions. A fitted trend line is presented in blue as the mean values +/-standard deviation. The standard deviation is shaded in light gray.

We also examined eQTL whose effects may vary as a function of genetic ancestry. Our examination followed a similar model to the main effect analysis but with an interaction term between SNP and ancestry proportion. We identified at least one ancestry-dependent cis-eQTL (within +/-500 kb of each feature) for 943 unique genes across brain regions (lfsr < 0.05, n=531, 942, 573, and 531 for the caudate, dentate gyrus, DLPFC, and hippocampus, respectively; **Fig. S22**, **Table S5**, and **Data S7**) with 54.1% (510 eGenes) shared across the four brain regions (**Fig. S23**). This relatively limited detection of ancestry-dependent eQTL supports other work showing high correlation of causal effects across local ancestry of admixed individuals (*32*).

We next tested whether these eGenes (main effect and ancestry-dependent) were likely to be differentially expressed by genetic ancestry. Across brain regions, we found significant enrichment (Fisher’s exact test, FDR < 0.05) of these eGenes (lfsr < 0.05) with ancestry-associated DEGs (lfsr < 0.05; **Fig. 4C** and **Fig. S23C**). Given the potential correlation of genotypes with eGenes and ancestry inference, we also examined allele frequency differences between DEGs and non-DEGs. We found a significant increase in allele frequency differences for DEGs compared with non-DEGs (Mann-Whitney U, p-value < 0.05; **Fig. 4D** and **Fig. S24**) across brain regions. These results suggest that a genetic component is likely influencing these expression differences, potentially due to divergence in allele frequencies.

To test this possibility, we imputed gene expression levels from genotypes using an elastic net model, and then examined the correlation between the observed genetic ancestry effect from our ancestry DE analysis and the predicted genetic ancestry effect computed from the predicted expression levels across samples. Unsurprisingly, eGenes showed higher prediction accuracy than non-eGenes; interestingly, however, eGenes with an ancestry difference in gene expression had a stronger genetic component (higher *R^2^*) than eGenes without an ancestry difference across the four brain regions (**Fig. S25**). Furthermore, the imputed gene expression levels explained an average of 59.5%, 58.7%, 56.8%, and 56.8% of the variance in genetic ancestry effect sizes across the caudate, dentate gyrus, DLPFC, and hippocampus, respectively (**Fig. 4E**). This variance explained generally increased on the isoform level (transcript [*R^2^* = 50.8%±7.0%], exon [*R^2^*= 61.6%±4.1%], and junction [*R^2^* = 62.6%±5.1%]; **Fig. S26**) across brain regions. In contrast, the genetic variant for the top main effect eQTL associated with these genes explained on average ∼20% of the variance in genetic ancestry effect sizes with a similar proportion for the isoform level (**Fig. S27**). Thus, genetic variants contributed to nearly 60% of the observed genetic ancestry in gene expression – and variant effects on alternative splicing were even greater.

### Differential gene expression in a binary contrast of Black and White Americans

To extend our analysis of DEGs driven by genetic ancestry, we performed a binary analysis of combined Black and White American samples (**Table S6**) – the latter showing very little admixture of African ancestry (STRUCTURE; African ancestry mean = 0.03, range = 0-0.16; **Fig. S1**). Using these American samples, we selected individuals with relatively limited admixture (Black Americans ≥ 0.8 African genetic ancestry and White Americans > 0.99 European genetic ancestry) across the caudate, dentate gyrus, DLPFC, and hippocampus. To limit the influence of the larger sample size for this binary analysis (Black American vs White American), we randomly sampled ten times without replacement to approximate the admixed Black American-only analysis sample size (caudate, n=122 [61 each]; dentate gyrus, n=46 [23 each]; DLPFC, n=124 [62 each]; and hippocampus, n=134 [67 each]). We identified more than double as many ancestry-associated DEGs (5,324 unique genes, median lfsr < 0.05; **Fig. S28A**, **Table S7**, and **Data S8**) representing 28% of all genes tested across the caudate (n=2,877), dentate gyrus (n=2,219), DLPFC (n=3,318), and hippocampus (n=2,818) with similar immune system enrichment patterns (**Fig. S28B** and **Data S9**).

We next compared the binary analysis DE results (genes, transcripts, exons, and junctions) with the admixed Black American-only results. While we found a significant overlap of ancestry associated DE features (Fisher’s exact test, p-value < 0.0001), approximately 72% of features (3847 unique genes) were unique to the binary DE results (**Fig. S29**). Even so, effect sizes from binary analysis were significantly correlated (Spearman, rho = 0.43 to 0.49, p-value < 0.0001; **Fig. S30**) with effect sizes from admixed Black American-only analysis across features and brain regions, which increased when we examined only shared features (Spearman, rho = 0.60 to 0.66, p-value < 0.0001; **Fig. S31**). While these results confirm most of the ancestry-associated DEGs in the Black American sample alone, they also highlight additional ancestry-related factors that influence gene expression presumably including environmental events (i.e., epigenetic).

### Environmental contributions to global ancestry-associated differential expression

Our binary DE analysis of Black and White Americans suggests that environmental factors may also contribute to global ancestry-associated DEGs. To identify DEGs driven by environmental factors, we used DNAm as an environmental proxy in Black Americans. We began by identifying the top 1% of variable CpGs that are likely driven by unknown environmental factors. We identified these CpGs by removing variation attributable to batch and to unknown technical and biological factors as captured by the top five DNAm principal components, while preserving variation due to global ancestry. We then grouped those top variable CpGs into variable methylated regions (VMRs) for the caudate (89 samples; 12,051 VMRs), DLPFC (69 samples; 9,701 VMRs), and hippocampus (69 samples; 9,924 VMRs). In contrast to our DE analysis results, we identified fewer VMRs that were differentially methylated regions (DMRs) for global ancestry (FDR < 0.05; n=3, 1, and 8 for the caudate, DLPFC, and hippocampus, respectively). However, we identified a larger number of local ancestry-associated DMRs (FDR < 0.05; n=494, 260, and 265 for the caudate, DLPFC, and hippocampus, respectively; **Fig. 5A**).

**Fig. 5:**
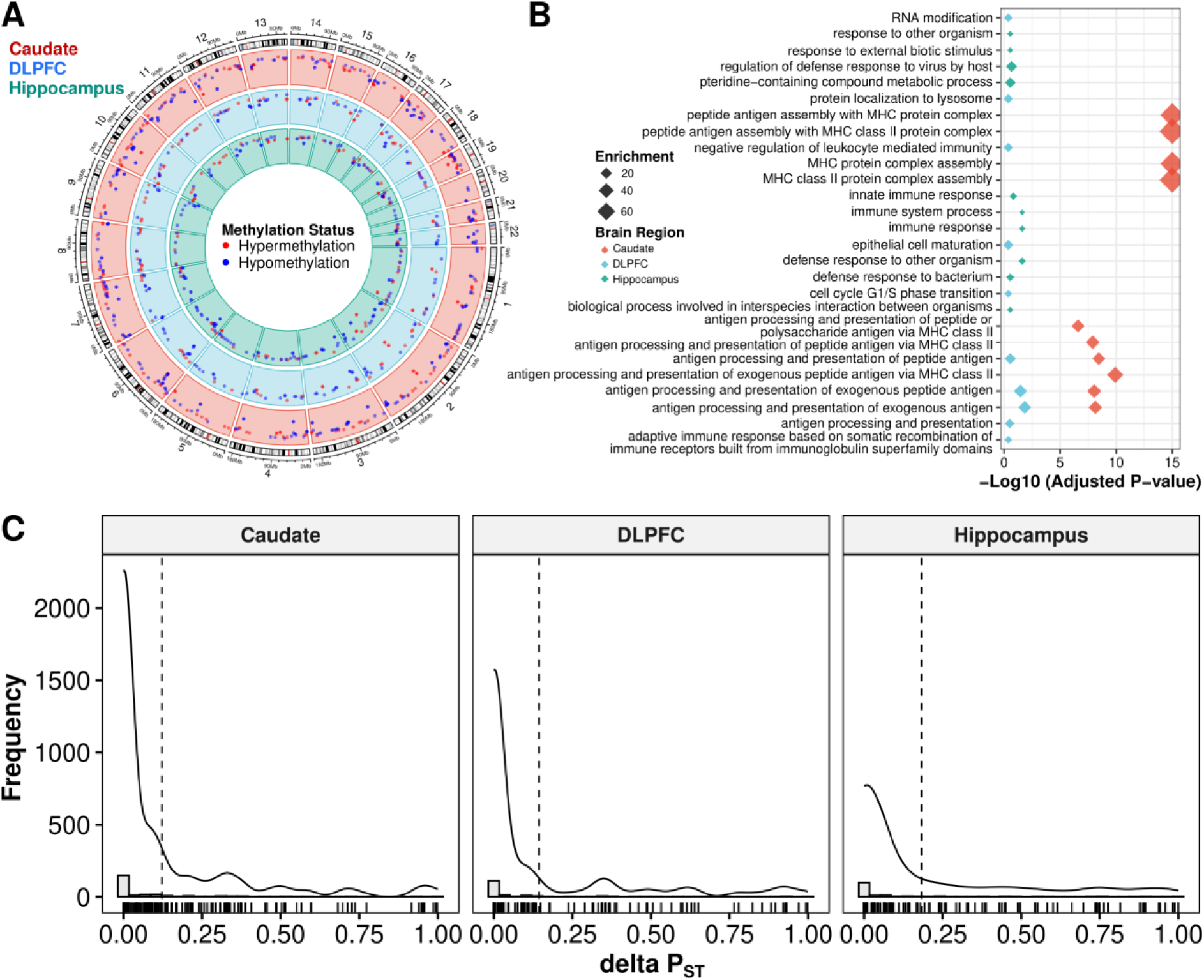
Unknown environmental factors are primary drivers of nearby global ancestry-associated DEGs. **A.** Circos plot showing local ancestry-associated DMRs across the caudate (red), DLPFC (blue), and hippocampus (green). Methylation status is annotated in red for hypermethylation and blue for hypomethylation. **B.** Gene term enrichment of DMRs across brain regions. **C.** Histograms showing distribution of ΔP_ST_ associated with the impact of unknown environmental factors as captured by residualized VMR (corrected by local ancestry, age, sex, and unknown biological factors captured by PCA) for nearby global ancestry-associated DEGs. A dashed line marks the mean ΔP_ST_. A solid line shows the density overlay.

We reasoned that the difference in DMRs linked to global and local ancestry can be explained both biologically and statistically. Biologically, DNAm tends to be more influenced by local genetic variations. Statistically, local ancestry is more variable than global ancestry, which results in a higher power to detect DNAm differences and a smaller standard deviation in estimated effect size. This is demonstrated in **Fig. S32** and **Data S12**, where we compared DNAm levels against local ancestry and global ancestry levels for VMRs associated with local ancestry. Even so, we find significant correlation between local and global ancestry-associated DMRs (**Fig. S33**). Functional enrichment analysis of local ancestry-associated DMRs suggested that these DMRs were enriched for gene sets related to immune functions across all three brain regions (**Fig. 5B**), consistent with the functional enrichment results of ancestry-associated DEGs.

We next regressed out known biological factors (local ancestry, age, sex), as well as the potential batch effects and other unknown biological factors captured by the top five principal components of DNAm levels for each VMR. We used P_ST_ estimates (*18*) to provide a measure of proportion of overall gene expression variance explained by between-population differences. P_ST_ values range from 0 to 1, where values close to 1 imply the majority of expression variance is due to differences between populations. We defined deltaP_ST_ (ΔP_ST_) as the difference between P_ST_ values before and after regressing out the effect of VMRs associated with each gene. Therefore, ΔP_ST_ quantifies the proportion of ancestry-associated DEGs that are likely due to environmental exposures. Using this method, we found that across brain regions the average ΔP_ST_ was 15% (12.2%, 14.4%, and 18.3% for the caudate, DLPFC, and hippocampus, respectively; **Fig. 5C**). Altogether, these results imply that unknown environmental exposures measured by DNAm provide a minor contribution to the observed, primarily immune-related expression differences in our Black American neurotypical sample.

### Association of ancestry-associated expression differences with immune-and brain-related traits

We reasoned that ancestry-associated DEGs may contain risk genes that explain susceptibility of brain-related illnesses based on ancestry. To explore this hypothesis, we conducted stratified LD score regression (S-LDSC) to assess the polygenic contributions of global ancestry-associated DEGs to 17 brain-related traits (e.g., ADHD, autism, BMI, depression, and schizophrenia) (*33*). As our ancestry-associated DEGs were enriched for gene sets related to immune functions, we included five immune-related traits as a positive control in our S-DLSC analysis. Overall, we observed that ancestry-associated DEGs were enriched for heritability of neurological disorders and immune-related traits but not for psychiatric disorders and behavioral traits (**Fig. 6**, **Fig. S34**, and **Data S10**). This also included limited enrichment of peripheral immune function (*34–36*) (Fisher’s exact test, FDR < 0.05; **Fig. S35**), which is consistent with our previous enrichment showing a greater association with brain immune cell types compared to non-brain immune cell types (**Fig. S12**).

**Fig. 6:**
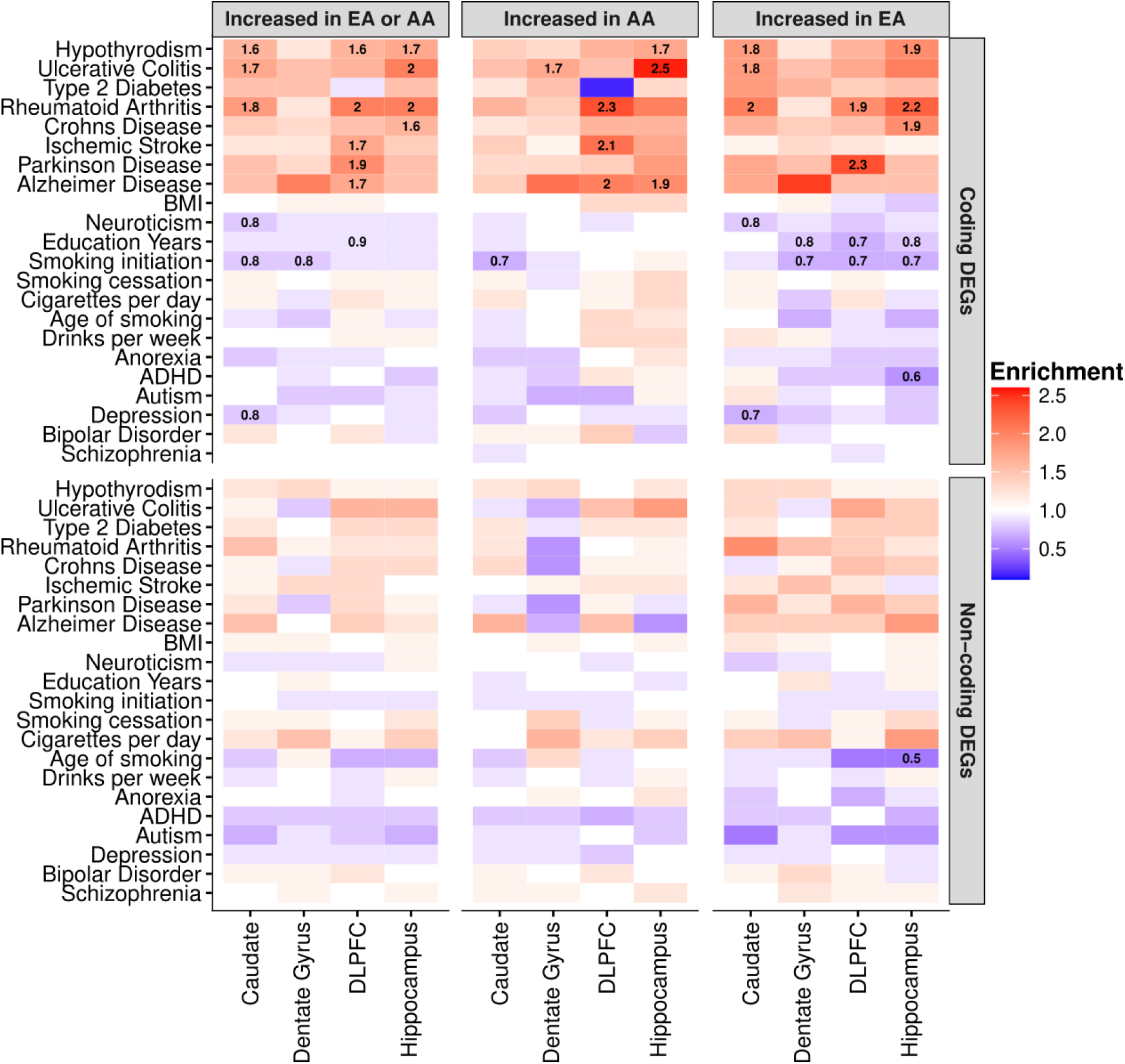
Global ancestry-associated DEGs stratified by coding or non-coding DEGs show general enrichment for heritability of several neurological and immune-related traits, but depleted for brain-related behavioral traits. Heatmap for ancestry-associated DEGs that show enrichment (red) or depletion (blue) for heritability of brain-and immune-related traits from S-LDSC analysis. Significant enrichment for heritability traits disappears when limited to non-coding DEGs. Numbers within tiles are levels of enrichment (> 1) or depletion (< 1) that are significant after multiple testing correction (FDR < 0.05). The left panel shows results for all DEG in each brain region. The middle and right panels show results for DEG increased with AA or EA proportions for each brain region, respectively.

Specifically, we found enrichment for heritability of ischemic stroke (enrichment fold = 1.5, FDR = 0.009) for ancestry-associated DEGs in the DLPFC, accounting for 26% of total heritability (**Fig. S34**). This enrichment was mainly driven by DEGs associated with an increase in AA proportion (enrichment fold = 1.7, FDR = 0.013), but not to EA (enrichment fold = 1.2, p-value = 0.2). Furthermore, stratified analysis by protein-coding and non-coding DEGs showed that enrichment was primarily driven by protein-coding DEGs, but not non-coding DEGs (**Fig. 6**). We observed stronger enrichment of ischemic stroke for protein-coding DEGs in the DLPFC (increased AA proportion; enrichment fold = 2.1, FDR = 0.011). This finding is consistent with epidemiological data that Black Americans are up to 50% more likely to experience ischemic stroke, and Black men are up to 70% more likely to die from stroke compared to non-Hispanic White men (*37*, *38*). Moreover, our cell-type enrichment analysis showed that the DEGs associated with increased AA proportion were enriched for vascular smooth muscle cells, endothelial cells, and pericytes (**Fig. S10**), all of which may contribute to vascular pathology implicated in stroke.

In addition to ischemic stroke, we also found enrichment for heritability of Parkinson’s disease (enrichment fold = 1.6, FDR = 0.025) for ancestry-associated DEGs in the DLPFC, accounting for 27% of disease heritability (**Fig. S34**). Interestingly, this enrichment was primarily driven by DEGs that were increased with EA proportion (enrichment fold = 1.9, FDR = 0.032), but not to AA proportion (enrichment fold = 1.3, p-value = 0.23). Again, this enrichment for Parkinson’s disease in the DLPFC was driven by protein-coding DEGs (increased EA proportion; enrichment fold = 2.3, FDR = 0.038; **Fig. 6**). This finding echos epidemiological studies suggesting that the prevalence of Parkinson’s disease is greater in White Americans compared with Black Americans (*39*). Additionally, cell-type enrichment analysis for DEGs associated with increased EA proportion showed enrichment for cell-type-specific genes related to the microglia, astrocytes, and oligodendrocyte progenitor cells (**Fig. S10**). Interestingly, we also found ancestry-associated glial cell subtypes (i.e., astrocyte [AST7] and oligodendrocyte lineage [OPC1]) significantly enriched for Parkinson’s disease heritability (enrichment fold > 2.0, FDR < 0.01; **Fig. S36**), suggesting a potential role for specific glial subtypes in the pathogenesis of Parkinson’s disease.

We also observed enrichment for heritability of Alzheimer’s disease for ancestry-associated DEGs across DLPFC, hippocampus and caudate accounting for 26%, 23% and 30% of total heritability, respectively (**Fig. S34**). These enrichments were mainly driven by protein-coding DEGs associated with an increase in AA proportion for the DLPFC (enrichment fold = 2.0, FDR = 0.013; **Fig. 6**) and hippocampus (enrichment fold = 1.9, FDR = 0.02; **Fig. 6**). Surprisingly, we found the opposite effect with an increase in EA proportion for the caudate when considering all DEGs (**Fig. S34**), which disappeared when considering only protein coding or non-protein coding DEGs (**Fig. 6**). Cell-type enrichment analysis of astrocytes, however, shows ancestry-specific effects consistent with this finding for the caudate (increased EA proportion; **Fig. S10**). Moreover, we found ancestry-associated glial cell subtypes (i.e., microglia [MG0] and astrocyte [AST1 and AST7]) significantly enriched for Alzheimer’s disease heritability (enrichment fold > 2.2, FDR < 0.01; **Fig. S36**) and ancestry-associated DEGs enrichment for multiple activated microglia states (*40*) (**Fig. S37A**). Interestingly, these microglia states were associated with mouse Alzhiemer’s disease-associated microglial genes and Alzhiemer’s disease GWAS signals (**Fig. S37B**). We also observed significant enrichment of ancestry-associated DEGs primarily with Alzheimer’s disease-related DEGs between early Alzheimer’s and late Alzheimer’s (late response; **Fig. S38**).

In contrast, we observed significant depletion in heritability for several brain-related traits (e.g., education years, smoking initiation, age of smoking, schizophrenia, and depression; enrichment fold < 1, FDR < 0.05; **Fig. 6**, **Fig. S34**, and **Data S10**) of our ancestry-associated DEGs across brain regions. These results are consistent with our observations that ancestry-associated DEGs are depleted for gene sets related to the neuronal functions that are believed to play major roles in psychiatric disorders and behavior traits.

## Discussion

Here we provide the first detailed characterization of the impact of genetic ancestry on expression and DNA methylation in the human brain. Using admixed Black American donors, we have identified thousands of genomic features (i.e., genes, transcripts, exons, and junctions) associated with genetic ancestry and demonstrated that these features are evolutionarily less constrained. Approximately 60% of these ancestry-associated DEGs are associated with genetic variations. Our data show consistent enrichment for immune response pathways for genetic ancestry-associated DEGs and consistent absence of ancestry associations with neuronal functions. Furthermore, we found similar trends when we examined local genetic ancestry. Even so, given expression heritability is dominated (i.e., about 70%) by many small trans effects (*41*, *42*), we have chosen to focus primarily on global genetic ancestry.

Interestingly, our findings show the direction of enrichment varies by brain region for immune-related pathways, increasing in relation to AA proportion in caudate and increasing in relation to EA proportion in the other regions. Because the specific genes in these immune function sets vary somewhat across regions, it is tempting to speculate on 1) how genetics and the environment sculpt variation in this regional biology and 2) whether the functional and behavioral impact of these ancestry-associated DEGs depends on the biology of particular brain regions. However, there is no simple “up or down” bias to the functional associations independent of brain region. For example, if AA proportion is a risk factor to immune response in the caudate, then by the same reasoning AA proportion would be a protector factor for immune response in the hippocampus and prefrontal cortex. We considered that differences in directionality across regions may reflect variation in cell composition as the caudate was the only brain region without a laminar architecture. However, laminar architecture in the brain has generally implicated neuronal biology (*43*), which was not the case here (i.e., enrichment of immune-related pathways). Notably, virtually all of our findings are more significant at the isoform level, implicating gene splicing and processing as a more incisive method for explaining the effect of ancestry on gene expression.

Among the more striking findings of our data is the enrichment of heritability for neurological brain illness among ancestry-associated DEGs. Small vessel stroke and ischemic stroke are up to 50% more frequent in Black Americans (*37*, *38*), and here we show that heritability for stroke was enriched among DEGs that were increased in proportion to AA in our admixed Black population. In contrast, heritability for Parkinson’s disease, which is more prevalent in White Americans (*39*), was enriched among DEGs in proportion to EA. Interestingly, we observed a nearly two-fold enrichment for heritability of Alzheimer’s disease among DEGs that were increased with AA proportion in DLPFC and hippocampus, regions cardinally involved in Alzheimer’s disease. This observation echoes the fact that Alzheimer’s disease is twice as prevalent in Black Americans (*44*, *45*). However, general enrichment of DEGs for Alzheimer’s disease in the caudate associated with an increase in EA proportion highlights the potential regional complexity of the disorder in the brain as caudate is not generally considered a site of Alzheimer’s disease pathology. Ancestral DEGs increase heritability for several immune disorders and traits but not specifically in relation to either ancestry across the brain. It is noteworthy that the DEGs are not linked with heritability of psychiatric disorders and related behavioral traits, perhaps consistent with genes associated with these traits being especially enriched in neurons, which were again, conspicuously lacking in DEGs based on ancestry.

In addition to our analysis of the admixed Black American population, we also performed a combined analysis with White Americans. As an internal validation, we found significant overlap between this and our Black American-only analyses (i.e., DE), but a dramatic increase in the extent of differentially expressed features. Additionally, this combined analysis (Black and White Americans) revealed similar enrichment of the immune response, again in analogous alternating directionality depending on brain region. While these results implicate environmental exposures that might reflect systematic differences between the two ancestral groups, disambiguating genetic from environmental factors in this context is challenging. We, therefore, chose to examine the environmental impact on our Black American-only global ancestry-associated DEGs. To this end, we identified thousands of VMRs across the brain in this context.

To highlight those VMRs likely enriched for environmental influence, we focused on the top 1% of VMRs and looked for ancestry-associated DMRs within these genomic regions. Consistent with DE analysis, we found that local ancestry DMRs were enriched for genomic regions related to immune functions. When we used VMRs as an environmental proxy to examine the effect of environmental exposures on the DEGs, we found they explained, on average, roughly 15% of population differences in gene expression. Although we used local ancestry to correct for genetic background, we cannot be sure that the variation captured via methylation is solely attributed to environmental factors or that methylation can capture all environmental factors. A limitation of this study is the lack of social determinants of health information, which could have directly measured specific environmental exposures instead of using DNAm as a proxy. Nevertheless, our analyses demonstrate the potential to limit the impact of potentially systematic environmental factors by leveraging admixture populations for genetic ancestry analyses.

This enrichment in immune-related pathways is not altogether unexpected: a previous study showed population differences in macrophages associated with the innate immune response to infection (*18*). Furthermore, it is well documented that genetic variation is an important contributor to immune variation (*46–48*) and immune cell function (*34–36*). This research is particularly important for neuropsychiatric disorders (including schizophrenia, autism spectrum disorder, and Alzheimer’s disease) where the immune system has been implicated (*49–51*). Many of these neuropsychiatric disorders also show a racial health disparity (*44*, *52–54*). As a result, we examined our enrichment of immune function in more detail. Interestingly, we found little evidence that the MHC region, HLA variation, or glial cell composition drove our identified immune-response pathway enrichment. Additionally, we found stronger enrichment of brain immune compared with peripheral immune cell types, suggesting the potential involvement of a brain-specific immune response of these DEGs. Altogether, our results provide a starting point for further investigation for potential therapeutic interventions involving the immune response – therapeutic interventions that could address these health disparities.

In summary, we provide a detailed examination of the genetic and environmental contributions to genetic ancestry transcriptional changes in the brain. We leveraged genetic diversity within admixture populations to limit environmental confounders, resulting in converging evidence of the immune response in genetic ancestry-associated transcriptional changes in the brain. The research we have provided here substantively furthers our understanding of the contribution of genetic ancestry in the brain, opening new avenues to the development of ancestry-aware therapeutics and paving the way for equitable, personalized medicine.

## Acknowledgments

We would like to extend our appreciation to the Offices of the Chief Medical Examiner of Washington DC, Northern Virginia, Kalamazoo Michigan, Santa Clara County, University of North Dakota, and Maryland for the provision of brain tissue used in this work. Additionally, we also extend our appreciation to the late Dr. Llewellyn B. Bigelow and members of the LIBD Neuropathology Section for their work in assembling and curating the clinical and demographic information and organizing the Human Brain Tissue Repository of the LIBD. Finally, we gratefully acknowledge the families that have donated this tissue to advance our understanding of psychiatric disorders.

The authors are indebted to many colleagues whose advice and suggestions were critical in this work, including Hailiang Huang, PhD, Kafui Dzirasa, MD, PhD, Ambroise Wonkam, MD, PhD, Yasmin Hurd, PhD, Amanda Brown, PhD, and select leaders of Black In Neuro. The African Ancestry Neuroscience Research Initiative is a collaboration between the LIBD, Morgan State University, Duke University, and members of the community led by Rev. Dr. Alvin C. Hathaway. We are also grateful to the administrative support of Jean Dubose, Yanet Raesu, and Gwenaëlle E. Thomas, PhD.

## Author Contributions

Conceptualization: KJMB, SH, and DRW; Methodology: KJMB, QC, NJE, LAH-M, JMS, ACMP, AEJ, SH, and DRW; Software: KJMB, QC, and SH; Formal Analysis: KJMB, QC, and SH; Investigation: JHS and TMH; Data Curation: KJMB, NJE, LC-T, GP, and JEK; Writing – Original Draft: KJMB, QC, SH, and DRW; Writing – Review & Editing: KJMB, QC, LAH-M, LC-T, ACMP, TMH, JEK, AEJ, SH, and DRW; Visualization: KJMB and SH; Supervision: KJMB, SH, and DRW; Project Administration: KJMB and SH; Funding Acquisition: KJMB, LC-T, SH, and DRW.

## Data availability

Publicly available BrainSeq Consortium total RNA DLPFC and hippocampus RangedSummarizedExperiment R Objects with processed counts are available at http://eqtl.brainseq.org/phase2/. Publicly available BrainSeq Consortium total RNA caudate RangedSummarizedExperiment R Objects with processed counts are available at http://erwinpaquolalab.libd.org/caudate_eqtl/. Publicly available dentate gyrus RangedSummarizedExperiment R Objects with processed counts and phenotype information are available at http://research.libd.org/dg_hippo_paper/data.html. Analysis-ready genotype data will be shared with researchers that obtain dbGaP access. FASTQ files are available for total RNA dentate gyrus, DLPFC, and hippocampus via Globus collections jhpce#bsp2-dlpfc and jhpce#bsp2-hippo available at https://research.libd.org/globus/. For the caudate, FASTQ files are available via dbGaP.

We used publicly available single cell datasets. Glial subpopulation single-cell data from the human postmortem hippocampus astrocyte, microglia, and oligodendrocyte lineage is available from UCSC cell browser (*“Human Hippocampus Lifespan”* collection). The human PBMCs single-cell data is available from Zenodo (10.5281/zenodo.4273999). Multiple human brain region single-cell datasets (i.e., DLPFC, hippocampus, nucleus accumbens, amygdala, and subgenual anterior cingulate cortex) are available by brain region from GitHub (https://github.com/LieberInstitute/10xPilot_snRNAseq-human). Human microglial state dynamics in Alzhiemer’s disease single-cell data is available from http://compbio.mit.edu/microglia_states/.

## Code availability

All code and Jupyter Notebooks are available through GitHub at https://github.com/LieberInstitute/aanri_phase1 with more detail (*55*).

